# Microbial contamination screening and interpretation for biological laboratory environments

**DOI:** 10.1101/439570

**Authors:** Xi Li, Xue Zhu, Wenjie Wang, Kang Ning

## Abstract

Advances in microbiome researches have led us to the realization that the composition of microbial communities of indoor environment is profoundly affected by the function of buildings, and in turn may bring detrimental effects to the indoor environment and the occupants. Thus investigation is warranted for a deeper understanding of the potential impact of the indoor microbial communities. Among these environments, the biological laboratories stand out because they are relatively clean and yet are highly susceptible to microbial contaminants. In this study, we assessed the microbial compositions of samples from the surfaces of various sites across different types of biological laboratories. We have qualitatively and quantitatively assessed these possible microbial contaminants, and found distinct differences in their microbial community composition. We also found that the type of laboratories has a larger influence than the sampling site in shaping the microbial community, in terms of both structure and richness. On the other hand, the public areas of the different types of laboratories share very similar sets of microbes. Tracing the main sources of these microbes, we identified both environmental and human factors that are important factors in shaping the diversity and dynamics of these possible microbial contaminations in biological laboratories. These possible microbial contaminants that we have identified will be helpful for people who aim to eliminate them from samples.

**Importance:** Microbial communities from biological laboratories might hamper the conduction of molecular biology experiments, yet these possible contaminations are not yet carefully investigated. In this work, a metagenomic approach has been applied to identify the possible microbial contaminants and their sources, from the surfaces of various sites across different types of biological laboratories. We have found distinct differences in their microbial community compositions. We have also identified the main sources of these microbes, as well as important factors in shaping the diversity and dynamics of these possible microbial contaminations. The identification and interpretation of these possible microbial contaminants in biological laboratories would be helpful for alleviate their potential detrimental effects.

## Introduction

Indoor environments are important since most of us spend his/her time indoor for the most part of his/her life[1]. The microbial communities of these environments are of particular interests; in-depth studies of environmental microbes in the last decade have shed light on the subtle effects they have on human health[2]. For example, a chronic exposure to some fungi can cause asthma, but early life exposure to various mold and its derivatives can protect children from allergic and autoimmune diseases[3]. A growing number of studies have helped us estimate the microbial diversity in various indoor environments, and revealed that microbial diversity is closely related to the geographic locations[4], weather conditions[5, 6], populations[7], functions[8], and internal ventilation conditions[9].

Ironically, the microbial compositions from indoor environment in various types of biological laboratories are less well-understood. While microbial contaminants generally exist in molecular biology laboratories[10], few studies have been dedicated to study their microbial compositions. Biological laboratory contamination screening is an important task. Once a site is contaminated during the sampling process or the experiment procedure, the contaminants of the reagent or the environmental microbes may proceed to affect other samples, leading to biases in the results. It would also be intriguing to examine the hypothesis that each laboratory has a relatively stable microbial contamination, determined by various factors including the research subjects (such as animals, plants or microbes), personal factors, as well as macroscopic environment. Each type of microbial composition can then be used to characterize its associated type of laboratories, and help simplify future studies.

There are several approaches in the identification and quantification of microbial contaminants. The most commonly used technique is based on PCR amplification and sequencing of the genes which encode small subunit ribosomal RNA (16S rRNA). The alternative is the metagenomics approach, which sequences the DNA of the entire microbial community as a whole. Compared to culture-based approaches, metagenomic approaches are better for identifying novel organisms with unknown growth conditions[11]. High-throughput sequencing allows metagenomic approach to obtain all the genome information of the community in one experiment, enabling us to study the complex molecular interactions among species.

However, there are several difficulties in our application of the metagenomic approach. First, significant amount of microbial contaminants may be introduced during sample preparation, especially when sample has low microbial biomass. Second, unlike other well-studied environments, there is no catalog for quick screening of possible microbial contaminations from biological laboratory. Hence, it is imperative for us to design methods that could accurately identify microbial contaminants, trace the pollution source, and uncover their potential adverse effects.

To work out these problems, we collected samples from surfaces of several important sites (lab outlet, platform and the major public areas) of three types of biological laboratories (animal, plant and microbe), screened and annotated the microbial contaminants, identified the difference between sampling sites/laboratories, as well as discovered the microbial biomarkers for different types of biological laboratories. We also identified possible sources of these microbes, as well as the possible effects they may have on their occupants.

## Results and Discussions

### Compositions of microbial communities from different laboratories and different sampling sites

We obtained 759,612 high-quality 16s rRNA sequences in total for 37 samples. 724,126 sequences were retained after quality filtering, and all samples have reached the saturation plateau for sequencing, indicating enough sequences for 16s rRNA profiling. Among all sequencing data, 432,092 sequences were from microbiology laboratory (ML), 137,575 from animal laboratory (AL) and 154,460 from plant laboratory (PL). Then all sequences were clustered into 1,234 Operational Taxonomic Units (OTUs) at 97% similarity threshold. In order to ensure enough sequencing depth, we generated the rarefaction curves for each sample. At around 1,800 sequences per sample, most rarefaction curves showed saturation, suggesting that the depth of samples sequencing covered enough extent of taxonomic diversity.

To compare the microbial composition of all microbial contaminant samples from the animal, plant and microbe laboratories, the taxonomies at phylum- and genus-level were illustrated (**Figure 1**). The microbial communities are composed mainly of 6 different bacterial phyla, including *Proteobacteria*, *Actinobacteria*, *Bacteroidetes*, *Cyanobacteria*, *Firmicutes* and *Fusobacteria*, with differentiated proportions in each sample. *Actinobacteria* was the most abundant phylum across all samples (**Figure 1a**). At the genus level, *Proteus*, *Prevotella*, *Chryseobacterium*, *Methylobacterium*, *Acinetobacter*, *Enterobacter*, *Micrococcus*, *Rhodococcus*, *Stenotrophomonas* and *Staphylococcus* were the dominant components (**Figure 1b**). The microbial communities from various sites at genus level were very diverse, even from the same type of laboratory.

**Figure 1.**
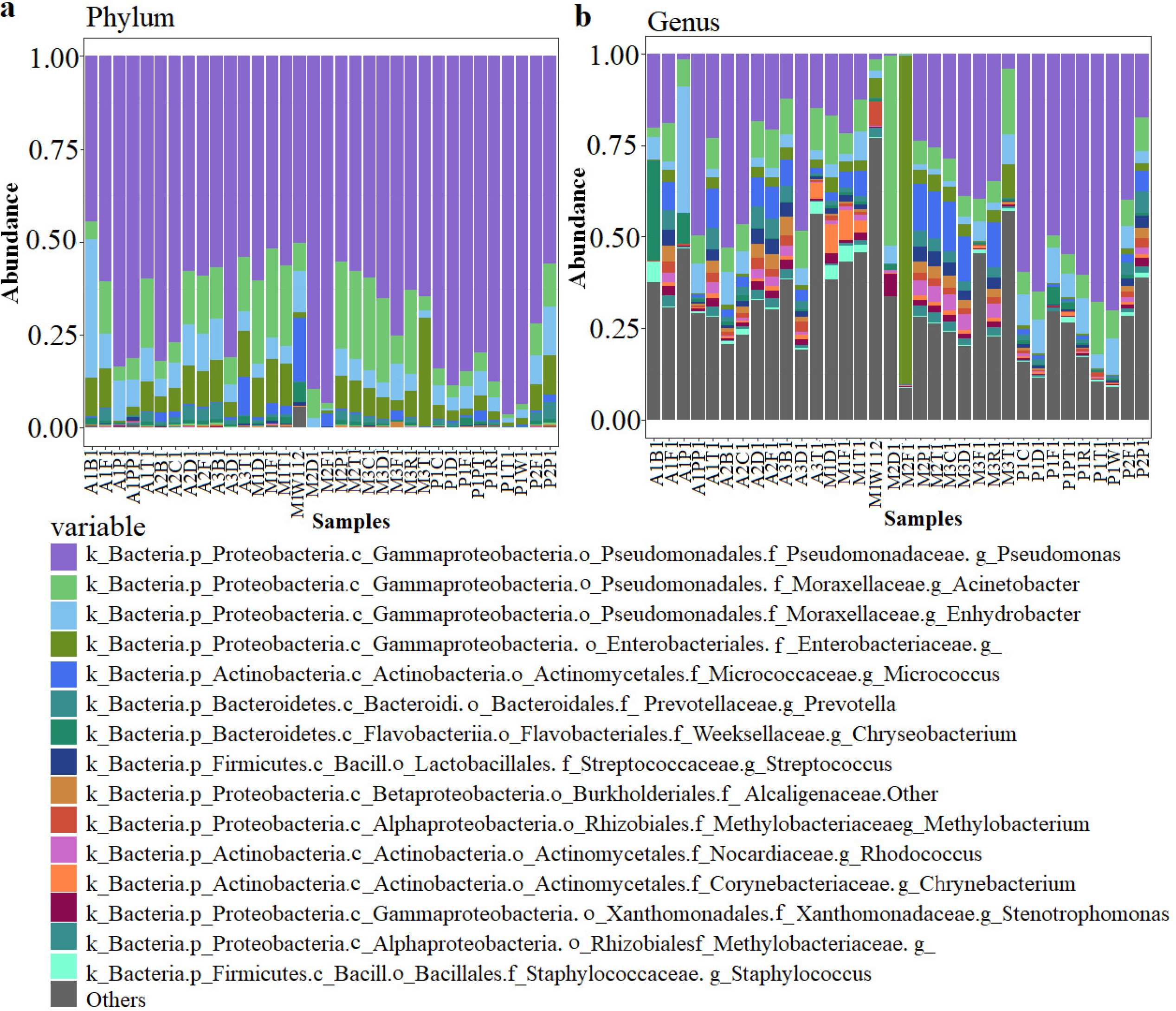
The relative abundance of the top 15 genera detected in samples across all laboratories. Each column represents a single sample, and sections **a** and **b** describe the same samples at different classification levels. **a**, At phylum level. **b**, At genus level. ‘Others’ indicates all other phyla or genera except for the top 15 genera.

### The relationship of microbial community composition, laboratory type and sampling sites

The type of laboratories carry more weight than sampling sites in the differentiation of microbial community samples. Alpha diversity analysis was performed (**Supplementary Table 1**), followed by the analysis of variance (ANOVA), to detect differences among samples from different sites and laboratories (**Figure 2**). Chaos indices showed that there is significant differences between AL and ML (**Figure 2a**). Shannon indices showedthat significant difference in the platform between ML and PL (**Figure 2b**). Furthermore, the number of OTUs determined by the Observed_OTUs revealed a clear difference from the major public areas between AL and PL (**Figure 2c**). **Figure 2** also shows that the samples from different types of laboratories could always be distinguished, whereas the samples within the same type of laboratories are usually indistinguishable except for the lab outlet and platform of PL. Thus, the differences in microbial community composition of samples across different types of laboratories are clear, while within laboratory differences are relatively small.

**Figure 2.**
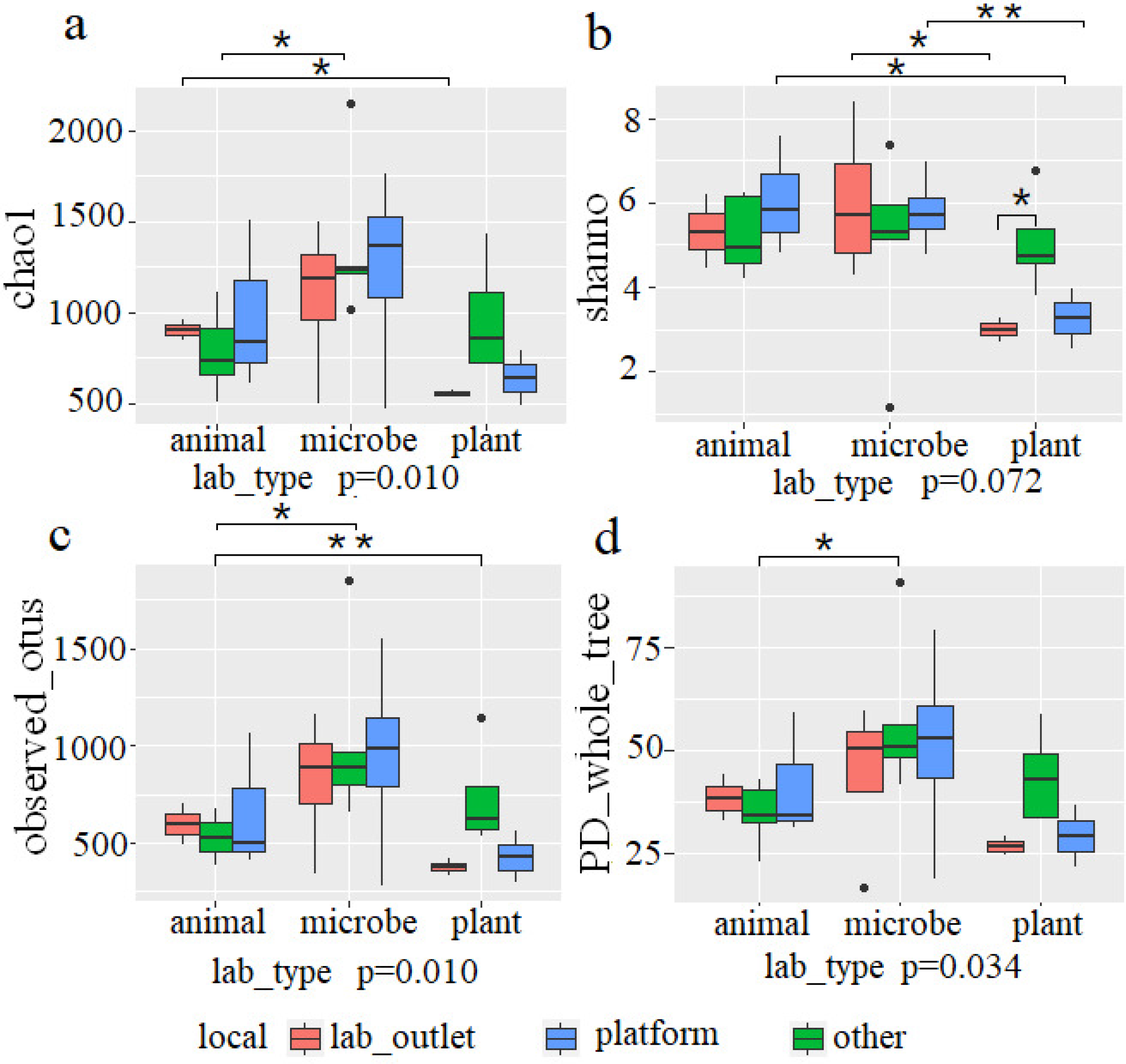
Alpha diversity comparisons for samples from all sampling sites/ biological laboratories. **a**, Chao1. **b**, Shannon index. **c**, Observed_OTUs. **d**, PD_whole_tree. Where Chao1 and Observed_OTUs estimate the number of OTUs in the community, and a higher Shannon index indicates greater abundance with a more even representation and PD_whole_tree adds the evolutionary relationship between species to compare its diversity. All samples have been compared with each other, categorized by different laboratories and sampling sites. The line indicates the difference between two sites, and *p < 0.1, **p < 0.01. ‘Other’ indicates the major public areas.

To gain further insights into the differences between laboratories, a comparison of samples from the same type sampling site across different types of biological laboratories was conducted. The results showed that these samples composed of many similar genus, but the proportion of each genus was different (**Figure 3**; **Table 1**). *Pseudomonas, Cinetobacter, Enterobacter* and *Micrococcal* were ubiquitous bacterial genus with dominant occurrence on the platform and lab outlet (**Figure 3a-c**). In addition, while the total number of detected genus are similar among lab outlet (76), public area (81) and platform (79), the number of shared genus is largest in public area (39), and smallest in lab outlet (22) (**Figure 3d-f**). Moreover, for either of the sampling sites including lab outlet, public area and platform, PL has much less laboratory-specific genus compared to AL and ML (**Figure 3d-f**). Therefore, we speculated that while public areas shared by experimenters might have largest number of shared genus, key sites such as lab outlet and platform has their specific sets of genus as potential contaminations.

**Figure 3.**
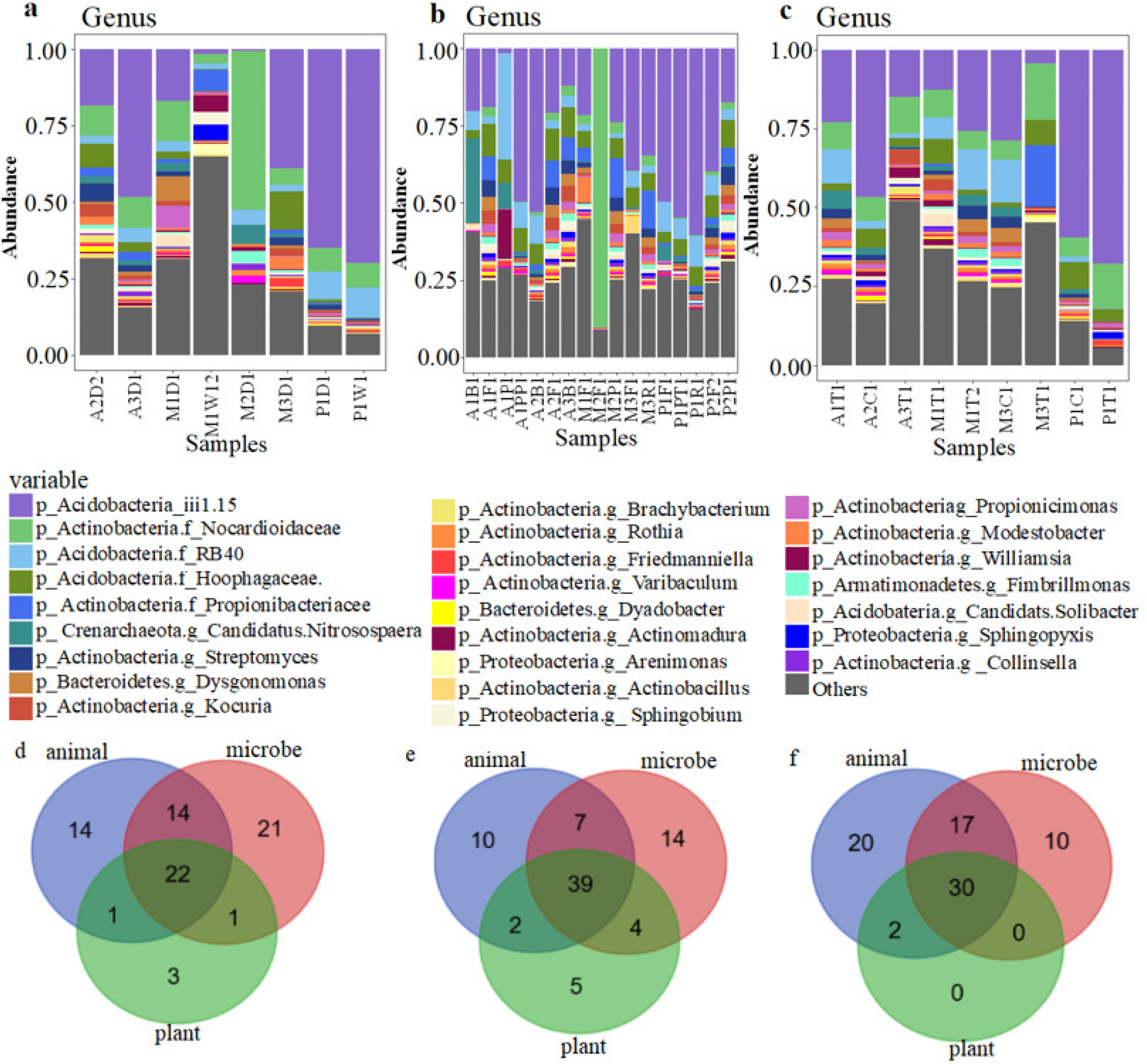
The composition of microbial samples at same types of sampling sites among all biological laboratories. The relative abundance of different species from the lab outlet (**a**), major public areas (**b**), and platform (**c**). Overlaps between the laboratories are indicated by Venn diagram showing the detected bacterial genera from lab outlet (**d**), major public areas (**e**), and platform (**f**).

**Table 1.** The relative abundance of shared genera across all laboratories and their possible sources. Although these bacteria exist in all laboratories, they exist in different proportions in each laboratory. Where ‘a’, ‘b’, ‘c’ represents the microbes that may be prevalently contaminated in laboratory reagents (a), introduced by human daily activities (b), and basic environmental microbe (c), respectively.

| Phylum | Genus | Relative abundance |  |  | Possible source |
| --- | --- | --- | --- | --- | --- |
|  |  | Animal laboratory | Microbiological laboratory | Plant laboratory |  |
| Actinobacteria | Corynebacterium | 0.01043194 | 0.020823612 | 0.005222726 | a, b, c |
|  | Brevibacterium | 0.004147394 | 0.00654011 | 0.001078781 | a |
|  | Micrococcus | 0.043620243 | 0.059833667 | 0.014597697 | a, b |
|  | Actinomyces | 0.00419055 | 0.011584026 | 0.001514026 |  |
|  | Kocuria | 0.002390392 | 0.004253708 | 0.001226562 | a |
|  | Propionibacterium | 0.003290537 | 0.010990419 | 0.008227309 | a, b |
|  | Rhodococcus | 0.013184083 | 0.019453764 | 0.004975993 | a |
|  | Microbacterium | 0.001041871 | 0.003651781 | 0.001070219 | a |
| Bacteroidetes | Sphingobacterium | 0.003109507 | 0.001733227 | 0.001112906 | b, c |
|  | Flavobacterium | 0.003071589 | 0.001690131 | 0.001168052 | a, b |
|  | [Prevotella] | 0.002920677 | 0.001850549 | 0.001664687 |  |
|  | Prevotella | 0.027960465 | 0.016342249 | 0.015979757 |  |
|  | Capnocytophaga | 0.004329217 | 0.001414413 | 0.004057175 |  |
|  | Chryseobacterium | 0.034069126 | 0.004402164 | 0.012406579 | a, b |
|  | Sediminibacterium | 0.006750533 | 0.002524042 | 0.003997523 |  |
|  | Porphyromonas | 0.001674581 | 0.001180926 | 0.001548046 |  |
| Firmicutes | Streptococcus | 0.020726981 | 0.017232988 | 0.009853175 | a, b, c |
|  | Bulleidia | 0.00260196 | 0.001167639 | 0.001166914 |  |
|  | Lactobacillus | 0.007177304 | 0.003449047 | 0.002745995 | b, c |
|  | Staphylococcus | 0.011641965 | 0.010651629 | 0.006485062 | b, c |
|  | Veillonella | 0.0060055 | 0.003390379 | 0.001033465 |  |
| Fusobacteria | Leptotrichia | 0.006702987 | 0.003816631 | 0.004186437 |  |
|  | Fusobacterium | 0.010868195 | 0.004798143 | 0.007385495 |  |
| Proteobacteria | Neisseria | 0.00857373 | 0.004033369 | 0.00663029 |  |
|  | Stenotrophomonas | 0.013283321 | 0.015911227 | 0.005654317 | a, b |
|  | Pseudoxanthomonas | 0.00734037 | 0.004651302 | 0.005937601 | a |
|  | Sphingomonas | 0.004221489 | 0.00850698 | 0.003254122 | a, b |
|  | Acinetobacter | 0.085566067 | 0.10519708 | 0.074663258 | a, b |
|  | Agrobacterium | 0.001620153 | 0.001257811 | 0.001000628 |  |
|  | Sphingobium | 0.006496551 | 0.003954288 | 0.004470411 | a, b |
|  | Paracoccus | 0.002275995 | 0.003590857 | 0.00553695 | a, c |
|  | Haemophilus | 0.004815064 | 0.002972872 | 0.001998964 |  |
|  | Methylobacterium | 0.016558476 | 0.014545819 | 0.008581459 | a, b, c |
|  | Pseudomonas | 0.272891893 | 0.191455024 | 0.538318946 | a, b, c |
|  | Brevundimonas | 0.006619573 | 0.009237963 | 0.002830279 | a, b, c |
|  | Enhydrobacter | 0.070222079 | 0.032529829 | 0.074842363 | a, b, c |
|  | Novosphingobium | 0.00188908 | 0.001385483 | 0.001607949 | a |
|  | Lysobacter | 0.001891199 | 0.002709745 | 0.001717886 |  |
| **Thermi** | Deinococcus | 0.008771836 | 0.008731487 | 0.005717905 |  |

We next compared the relative abundances of representative genus from three main sites within the laboratory. *Pseudomonas*, *Acinetobacter* and *Enterobacter* were most abundant among all sampling sites (**Figure 4a-c**). In addition, the number of all identified genus in AL (84) and ML (87) were much more than those in PL (53) (**Figure 4d-f**). Moreover, the platform of AL has the highest number of site-specific genus (**Figure 4d-f**). These results again confirm that the richness of microbial communities of platform and lab outlet depended heavily on the type of laboratory.

**Figure 4.**
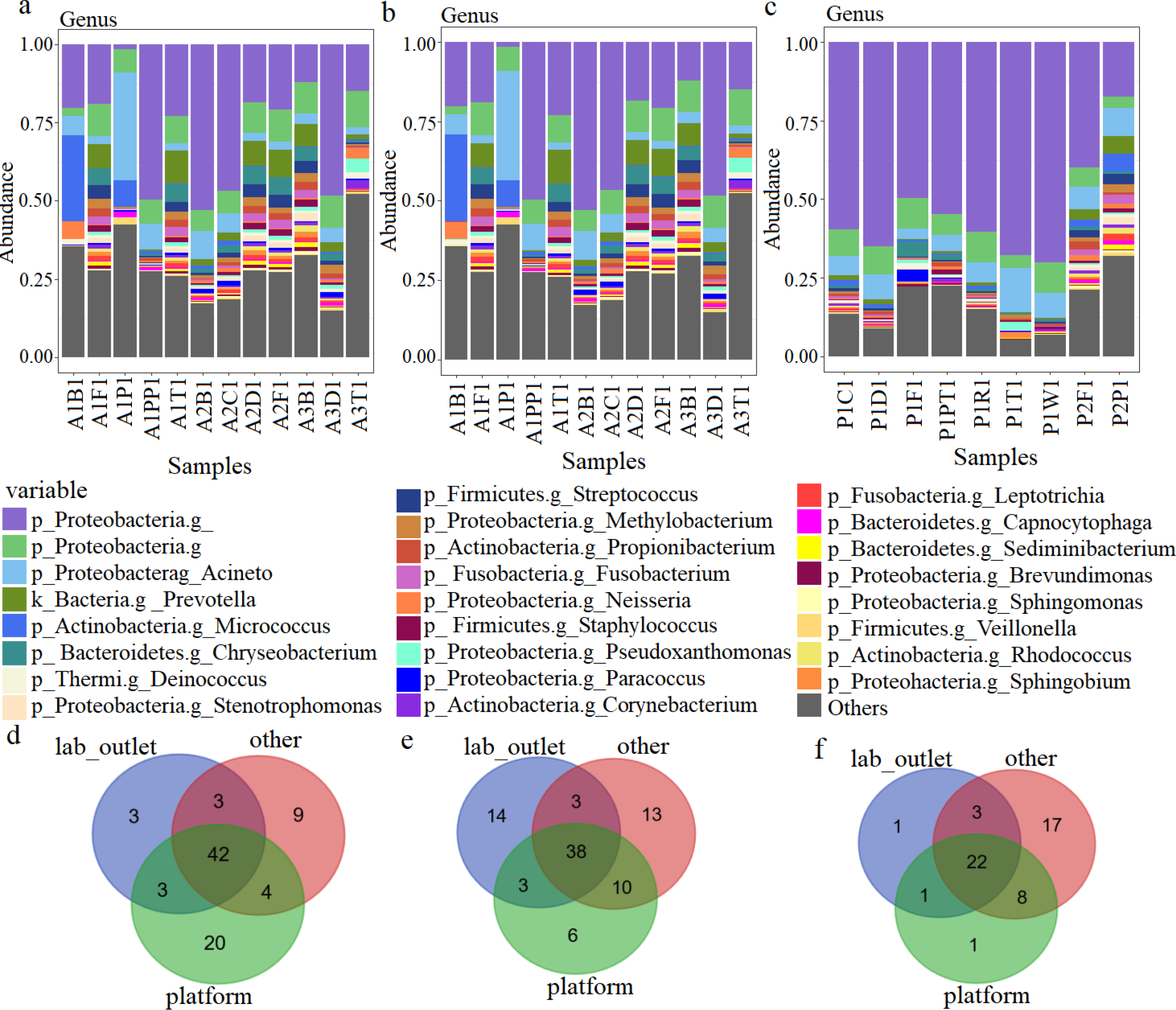
Differences of mirobial samples at different sites within the same type of biological laboratory. The relative abundance of different species from animal laboratory (**a**), microbiology laboratory (**b**) and plant laboratory (**c**). Venn diagram showing the overlap between identified microbial genera observed in animal laboratory (**d**), microbiology laboratory (**e**), plant laboratory (**f**). Where colors in **a**, **b**, **c** indicate various microbial genera, while ‘other’ in **d**, **e**, **f** indicates the major public areas.

### Possible sources and microbial biomarkers for different types of laboratories

We then performed literature mining to identify the possible sources of these microbial contaminations, referencing varies sources. We categorized the sources into laboratory reagent microbe, human-introduced microbe, and basic environmental microbe. Interestingly, laboratory reagents and human daily activities might play very important roles in introducing these possible microbial contaminations (**Table 1**).

To obtain a characteristic set of microbial contaminants, or biomarkers, for each type of biological laboratory, we used LDA Effect Size (LEfSe) to discover the biomarkers at each taxonomic level. 29 taxa (7, 15 and 7 taxa from AL, ML and PL respectively; **Figure 5**) were detected with high LDA scores. For samples from AL, *Becateroidetes*, *Flavobacteriaceae* and *Gemmata* were identified as biomarkers. *Enterobacteriales* and *Enterobacteriaceae* were identified as biomarkers for ML. *Pseudomonas*, *Pseudomonadaceae and Pseudomonadales*, which belong to *Pseudomonas,* were identified as biomarker for PL with high confidence (**Figure 5a**). The evolutionary relationship between these bacteria at different taxonomic levels is shown in **Figure 5b**.

**Figure 5.**
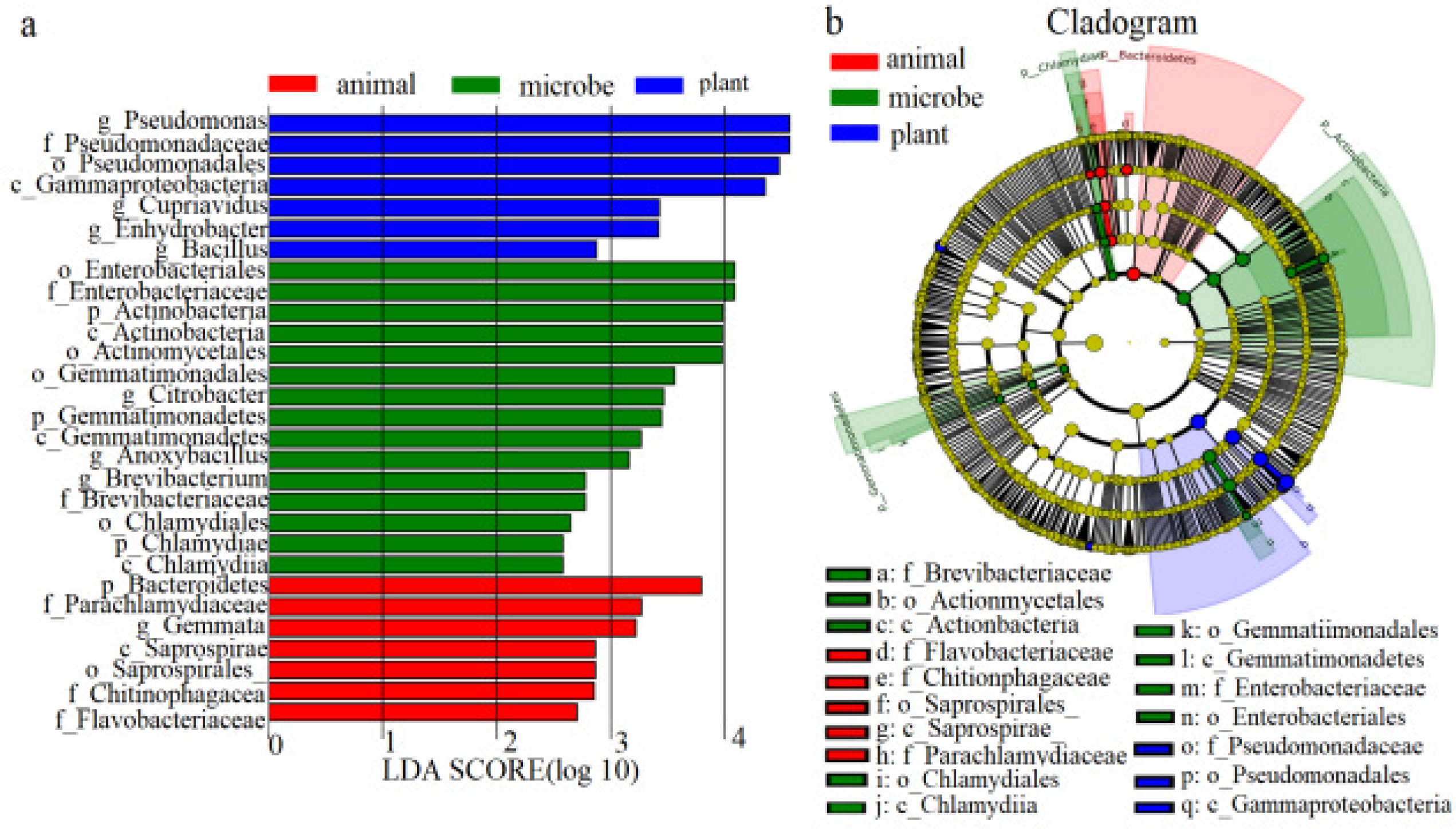
Biomarker for samples among three types of biological laboratories. **a**, Microbial richness that has significant differences between three types of laboratories (LDA > 2). **b**, The phylogenetic relationships of these microbes.

To further explore the characteristics of the biomarkers for different laboratories, we screened the genera with a relative abundance of > 1/1000 within the same type of laboratory. This identified ML to contain a greater variety of bacteria (65 genera) than AL (59) and PL (48). The population of the overlap between the detected genera of the three types of laboratories was 39 (**Table 1**), the highest was found between ALs/MLs (9 shared genera) and followed by ALs/PLs (3) and MLs/PLs (1), 16 specific genera in MLs, more than ALs (8) and PLs (5). Comparing against the references tables including reagents[12] (**SupplementaryTable 2**), residential[13] (including daily residential areas, office and classroom; **Supplementary Table 3**) and detected in ICU[14] contamination table (**Supplementary Table 4**), the shared genera exhibited significant overlap, while the specific genera did not. For laboratory-specific genera, only *Methyloversatilis* and *Psychrobacter* from AL was detected in reagent (representing laboratory reagent microbes) and ICU contamination table (representing basic environmental microbes) as mentioned above, while Bacillus from PL was observed in three reference tables, and *Flavisolibacter* from ML was only present in ICU contamination table. Therefore, we speculated that the overlapping specific- and shared- genera should be ubiquitous bacteria in the environment, lab reagents contaminants or external bacteria introduced by human activities.

Through literature mining, we assessed the possible effects of laboratory-specific genera (**Table 2**) without any overlap with the three reference tables. Specific bacteria of laboratory will assert adverse effects on researchers or experimental materials. To illustrate, the *Jeotgalicoccus* of AL as a pathogen, can be transmitted via air or surfaces contact and hence infects hosts; *Moraxella* of PL could influence the onset of bronchitis or pneumonia. Other microbes are less harmful; for instance, *Psychrobacte* of AL is a probiotic of fish, and its highest diversity was detected in sample A1B1, corresponding to the incubator of the zebrafish laboratory by backtracking analysis. *Buchnera* of PL, a symbiotic bacterium of aphids is specifically associated with the tissue culture process. *Flavisolibacter* of ML, which improves nitrogen fixation in rhizosphere of plants, has the highest abundance in sample M1W12, which was from cultivated plants on the windowsill in the M1 laboratory. Together, these results showed high concordance between the characteristics of the laboratory and the sampling site, demonstrating that the compositions of microbial communities have profound association with their hosting laboratories.

**Table 2.**
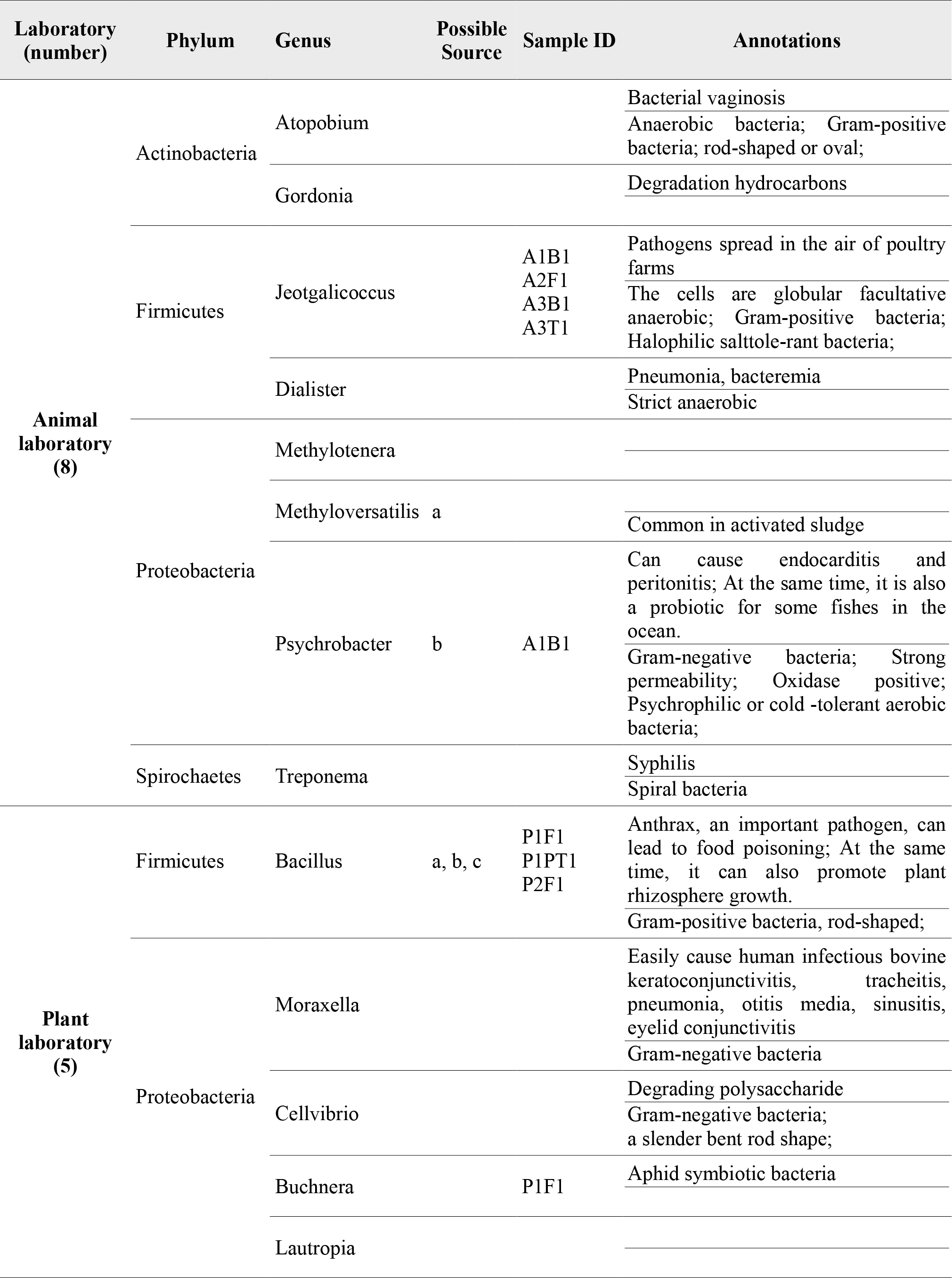
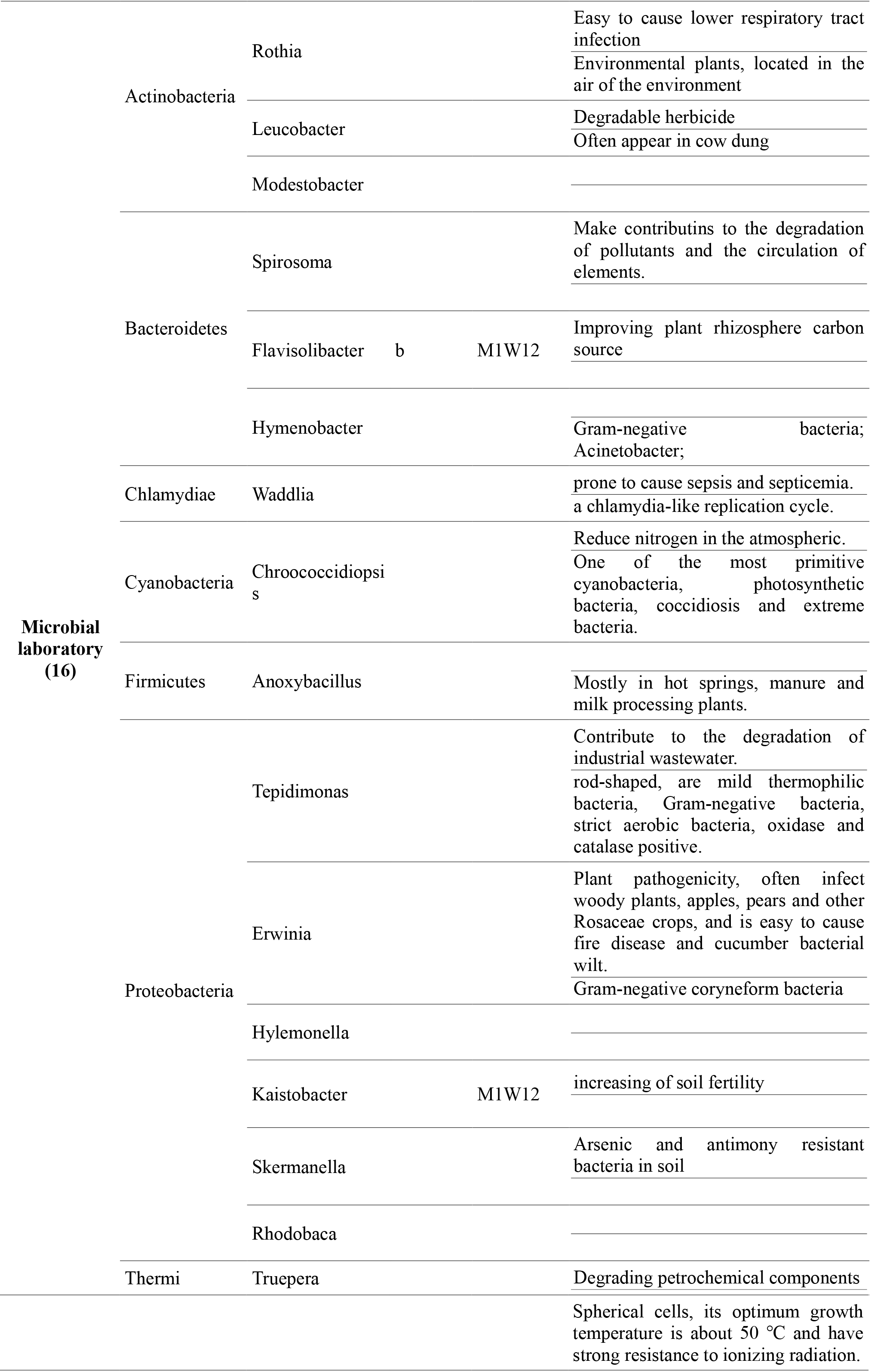
The specific genera of the three types of biological laboratories. We labeled the basic features of these bacteria, identified these potential effects through literature mining, and marked the samples in which the species were most abundant for subsequent studies.

As already known, the present of these contaminants can bring inconvenient for our experiment more or less, so caution and preciseness must be followed throughout the whole experiment. And the use of blank control during sampling, DNA extraction and sequencing is also necessary for detecting contamination. Furthermore, he contaminants are associated with the use of different kits or batches, which can introduce variation in reagent contamination[12], therefore, it would be best to use the same kits in one experiment and disentangle batch effects. Additionally, we should catalogue the laboratory microbial contaminants better, and thus, as if we know the contaminants, antibiotic treatment can be executed before experiment to mitigate the experimental bias caused by these microbial contaminants.

## Conclusion

In this work, a metagenomic approach has been applied to identify the possible microbial contaminants and their sources, from the surfaces of various sites across different types of biological laboratories. The possible microbial contaminants that we have identified will be helpful for people who aim to eliminate them from samples.

As far as we know, our work is the first investigation on the composition of microbial communities in biological laboratories. We found several interesting patterns in these compositions. First, there are significant differences in the structures of the microbial communities from the three types of laboratories. Factors such as sampling sites (including lab outlet, platform and the major public areas) and laboratory types (for animal, plant and microbe), have influenced the compositions of indoor microbial communities: the number of microbial genus in animal and microbial laboratories are significantly higher than those in plant laboratories, while key sites such as lab outlet and platform have their specific sets of genus as potential contaminations for each type of laboratory. These differences are highly related to the functions of the laboratories. Second, the type of laboratories has more influence than sampling sites in the differentiation of microbial community samples. Third, while public areas shared by experimenters may have the largest number of shared genus, key sites such as lab outlet and platform have their specific sets of genus as potential contaminations for each type of laboratory. This suggests that while general human activities have the most effect on the microbial community structure of the laboratory, the microbial communities of platform and lab outlet depends more heavily on the type of the laboratory. Finally, by tracking the possible sources of laboratory microbes, we found that laboratory reagents and human daily activities might play very important roles in introducing these possible microbial contaminations. These microbes are intimately connected with the experimental materials, and will also assert negative effects on the experiment process as well as on experimenters.

We would like to suggest two directions in future analysis of possible microbial contaminations from laboratories. First, better profiles of the microbial compositions in the biological laboratories are needed. They would help in devising countermeasures to mitigate the experimental bias caused by these microbial contaminants. Second, we hope that longitudinal studies would help to confirm our findings, since our samples were collected from the same building in summer and may not reflect the seasonal dynamics of the microbial communities.

## Materials and Methods

### Experimental design and sample collection

We selected 8 laboratories (3 animal laboratories, 3 microbial laboratories, 2 plant laboratories) from the College of Life Science and Technology, Huazhong University of Science and Technology in Wuhan, Hubei province of China. All laboratories are in the same building, but distributed at different floors. We collected samples from the lab outlet (e.g. doors, windows) with high air mobility, the platform (e.g. processing table, clean bench), and the major public areas (e.g. floor, pool and preprocessing pond). We conducted all sampling and genome extraction in July of 2017 to avoid the influence of environmental and climate factors. The overall experimental design, and main methods of our project are shown in **Figure 6**.

**Figure 6.**
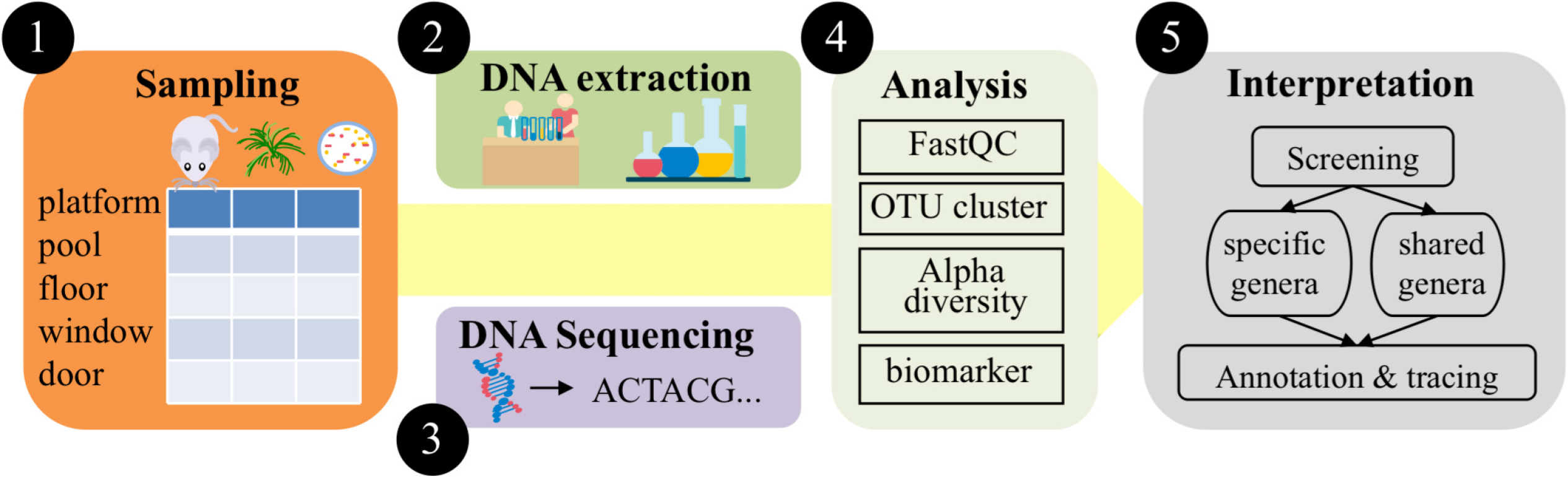
Schematic workflow including sampling site selection, DNA sequencing and computational approaches. Illustration of the main steps involved in sampling from lab outlet, platform and major public areas of animal, microbe and plant laboratories, extracting DNA, Illumina sequencing, bioinformatics analysis and interpretation. Finally, we compared the detected genera with the publicly available common contaminants in the reagent, ICU microbe table and the basic microbes of the environment to annotate the bacteria and trace the possible pollution source.

All samples were wiped on the selected surface areas and devices with 4 to 5 swabs that were moistened with a 15 mL centrifugal tube containing 2.5 mL of normal saline. All sampling locations, primer used and their characteristics are listed in **Supplementary Table 5**. During sampling, all the staff and devices were in full operation (normal status). After sampling, the swabs were kept at 4 °C. Afterwards, the genome of all samples extracted by using biological sampling kit was stored at −20 °C.

### DNA extraction

First, 1 mL sample, in total, and 1 mL buffer was added into centrifuge tube, and the mixture was stirred gently. After water bath of 2 h 65 °C, mixing by hand every 30 min, the suspension was vortexed for 10 seconds. The tube was placed on ice for 10 min and centrifuged afterwards (100 g, 5 min, 4 °C). The supernatant was transferred into another tube, and an equal volume of phenol: chloroform: isoamyl alcohol in a ratio of 25:24:1 was added. The suspension was mixed gently and centrifuged at 15°C/1000g for 5 min. The aqueous layer was transferred into a new tube. Then, the same volume of isopropanolis was added to cause the DNA to precipitate out of the aqueous solution. After incubation at −20 °C overnight, the suspension was centrifuged at 4°C/13500g for 30 min. After removing the supernatant, the precipitate was rinsed with 1 mL of 70% ethanol, and centrifuged repeated at 4 °C/13500 g for 30 min until the precipitate was completely dried and re-dissolved in 20 μL of PCR-grade water for easy handling and storage.

### 16S rRNA gene amplification and Illumina sequencing

For Illumina sequencing, 16 rRNA gene was amplified in the PCR reaction mixture (20 μl), which contained 1 μl Taq polymerase, 0.25 μl of forward primer, 0.25 μl of reverse primer, 0.5 μl of Dntp, 1 μl of template DNA, 5 μl 10×buffer and 12 μl ddH_2_O. To reduce the nonspecific amplification, the PCR system was made up on the ice box. The amplification process is as follows: 95 °C for 5 min, 25 cycles of 94 °C for 30 s, 54 °C for 40 s, 72 °C for 1 min, then followed by 30 cycle of 72 °C for 10 min and 4 °C hold. Amplification products were visualized with e gel. After quality filtering, the products was purified using the kits, and restored at −20 °C, then sent to company for Illumina sequencing. All sequencing data are deposited to NCBI SRA with project accession number PRJNA490598.

### Bioinformatics and statistical analysis of sequencing results

After obtaining the sequencing data for these samples, we used FastQC(http://www.bioinformatics.babraham.ac.uk/projects/fastqc/)[15] to perform preliminary quality control and filtering of the data.

QIIME (Quantitative Insights Into Microbial Ecology; http://qiime.org/)[16] is used for 16S rRNA profiling. Using the QIIME script join_paired_ends.py to process the double-ended sequences, merge them, and make the Mappingfile containing SampleID, BarcodeSequence, LinkerPrimerSequence, ReversePrimer, Description information. Then, we used validate_mapping_file.py to check the Mappingfile, and marked the wrong locations in the finally Mappingfile.html. Based on the extracted barcode information by referencing the Mappingfile with extract_barcodes.py, we split the sample by split_libraries_fastq.py, where the quality threshold was set to 20 (99% accuracy), then removed chimeras and length-marginized sequences.

Four common alpha diversity metrics and pick_de_novo_otus.py were used to extract OTUs from the Fasta file, removed the single reads from OUTs and obtained the rarefaction curve of the sample to determine the depth of the sequencing by filter_otus_from_otu_table.py and alpha_rarefaction.py. Biome summarize-table for counting the number, average number, and total number of sequences contained in each sample, alpha_diversity.py and beta_diversity_through_plots.py for analyzing the diversity of samples. Statistical analysis and visualization were then performed in R (https://www.r-project.org/)[17] using the package ggplot2. We then used SPSS (https://www.ibm.com/analytics/datascience-/predictive-analytics/spss)[18] to perform ANOVA on the alpha diversity results of samples to compare the difference of microbial community composition among the three sites.

LEfSe (LDA Effect Size; http://huttenhower.sph.harvard.edu/galaxy/) is used to find the biomarkers in the sample. The input file was obtained by summarize_taxa.py. In each group, the biomarker, LDA values and the hierarchical relationship between individual biomarkers were shown by run_lefse.py, plot_cladogram.py and plot_features.py severally.

## Acknowledgments

This work was partially supported by National Science Foundation of China grant 31871334 and 31671374, and Ministry of Science and Technology’s grant 2014AA021502 and 2018YFC0910502.

## References

1. Kelley ST, Gilbert JA: Studying the microbiology of the indoor environment. Genome Biology 2013, 14(2):1–9.

2. O’Hara NB, Reed HJ, Afshinnekoo E, Harvin D, Caplan N, Rosen G, Frye B, Woloszynek S, Ounit R, Levy S: Metagenomic characterization of ambulances across the USA. Microbiome 2017, 5(1):125.

3. Rook GA: Review series on helminths, immune modulation and the hygiene hypothesis: the broader implications of the hygiene hypothesis. Insect Science 2010, 126(1):3–11.

4. Chase J, Fouquier J, Zare M, Sonderegger DL, Knight R, Kelley ST, Siegel J, Caporaso JG: Geography and Location Are the Primary Drivers of Office Microbiome Composition. Msystems 2016, 1(2):e00022–00016.

5. Proctor CR, Dai D, Edwards MA, Pruden A: Interactive effects of temperature, organic carbon, and pipe material on microbiota composition and Legionella pneumophila in hot water plumbing systems. Microbiome 2017, 5(1):130.

6. Rintala H, Pitkäranta M, Toivola M, Paulin L, Nevalainen A: Diversity and seasonal dynamics of bacterial community in indoor environment. Bmc Microbiology 2008, 8(1):56.

7. Meadow JF, Altrichter AE, Kembel SW, Kline J, Mhuireach G, Moriyama M, Northcutt D, O’Connor TK, Womack AM, Brown GZ: Indoor airborne bacterial communities are influenced by ventilation, occupancy, and outdoor air source. Indoor Air 2014, 24(1):41–48.

8. Korpi A, Pasanen AL, Pasanen P: Volatile compounds originating from mixed microbial cultures on building materials under various humidity conditions. Appl Environ Microbiol 1998, 64(8):2914–2919.

9. Gilbert JA, Stephens B: Microbiology of the built environment. Nat Rev Microbiol 2018.

10. Salter SJ, Cox MJ, Turek EM, Calus ST, Cookson WO, Moffatt MF, Turner P, Parkhill J, Loman NJ, Walker AW: Reagent and laboratory contamination can critically impact sequence-based microbiome analyses. Bmc Biology 2014, 12(1):87.

11. Hugenholtz P, Goebel BM, Pace NR: Impact of culture-independent studies on the emerging phylogenetic view of bacterial diversity. Journal of Bacteriology 1998, 180(18):4765–4774.

12. Goffau MCD, Lager S, Salter SJ, Wagner J, Kronbichler A, Charnockjones DS, Peacock SJ, Smith GCS, Parkhill J: Recognizing the reagent microbiome. Nature Microbiology 2018.

13. Meadow JF, Altrichter AE, Kembel SW, Moriyama M, O’Connor TK, Womack AM, Brown GZ, Green JL, Bohannan BJM: Bacterial communities on classroom surfaces vary with human contact. Microbiome 2014, 2(1):7.

14. Oberauner L, Zachow C, Lackner S, Högenauer C, Smolle KH, Berg G: The ignored diversity: complex bacterial communities in intensive care units revealed by 16S pyrosequencing. Sci Rep 2013, 3(3):1413.

15. Brown J, Pirrung M, Mccue LA: FQC Dashboard: integrates FastQC results into a web-based, interactive, and extensible FASTQ quality control tool. Bioinformatics 2017, 33(19).

16. Lawley B, Tannock GW: Analysis of 16S rRNA Gene Amplicon Sequences Using the QIIME Software Package. Methods in Molecular Biology 2017, 1537:153.

17. Sunil Bhavsar PD, Shantilal Tank: R software package based statistical optimization of process components to simultaneously enhance the bacterial growth, laccase production and textile dye decolorization with cytotoxicity study. Plos One 2018, 13(5):e0195795.

18. Gouda MA: Common Pitfalls in Reporting the Use of SPSS Software. Medical Principles & Practice 2015, 24(3):300–300.

